# Primary infection with Zika virus provides one-way heterologous protection against Spondweni virus infection in rhesus macaques

**DOI:** 10.1101/2022.12.16.520792

**Authors:** Anna S. Jaeger, Chelsea M. Crooks, Andrea M. Weiler, Mason I. Bliss, Sierra Rybarczyk, Alex Richardson, Morgan Einwalter, Eric Peterson, Saverio Capuano, Alison Barkhymer, Jordan T. Becker, Joseph T. Greene, Tanya S. Freedman, Ryan A. Langlois, Thomas C. Friedrich, Matthew T. Aliota

**Affiliations:** Department of Veterinary and Biomedical Sciences, University of Minnesota, Twin Cities; Department of Pathobiological Sciences, University of Wisconsin-Madison; Wisconsin National Primate Research Center, University of Wisconsin-Madison; Department of Microbiology and Immunology, University of Minnesota, Twin Cities; Department of Biochemistry, Molecular Biology and Biophysics, University of Minnesota, Twin Cities; Department of Pharmacology, University of Minnesota, Twin Cities; Center for Immunology, University of Minnesota, Twin Cities; Masonic Cancer Center, University of Minnesota, Twin Cities

## Abstract

Spondweni virus (SPONV) is the closest known relative of Zika virus (ZIKV). SPONV pathogenesis resembles that of ZIKV in pregnant mice, and both viruses are transmitted by *Aedes aegypti* mosquitoes. We aimed to develop a translational model to further understand SPONV transmission and pathogenesis. We found that cynomolgus macaques (*Macaca fascicularis*) inoculated with ZIKV or SPONV were susceptible to ZIKV, but resistant to SPONV infection. In contrast, rhesus macaques (*Macaca mulatta*) supported productive infection with both ZIKV and SPONV and developed robust neutralizing antibody responses. Crossover serial challenge in rhesus macaques revealed that SPONV immunity did not protect against ZIKV infection, whereas ZIKV immunity was fully protective against SPONV infection. These findings establish a viable model for future investigation into SPONV pathogenesis, and suggest the risk of SPONV emergence is low in areas with high ZIKV seroprevalence due to one-way cross-protection between ZIKV and SPONV.

**Teaser:** Identification of asymmetric immune interactions between Zika and Spondweni viruses in macaque monkeys.

## Introduction

Arthropod-borne viruses (arboviruses) are increasingly contributing to the burden of human disease, and the mosquito-borne flaviviruses have caused significant epidemics during the past seven decades. Examples include the unprecedented rise in dengue virus (DENV) infections since World War II, the introduction of West Nile virus into the continental United States in 1999, the Zika virus (ZIKV) outbreak in the South Pacific in 2013-2014 and the explosive outbreak in the Americas in 2015-2016, ongoing yellow fever virus (YFV) outbreaks in Africa and Brazil, and the Japanese encephalitis virus outbreak in Australia in 2022. Although we cannot predict what might be coming next or when, RNA arboviruses can emerge unexpectedly to cause human disease on a global scale. The genus *Flavivirus* currently consists of ∼80 single-strand positive-sense RNA viruses (*1*), and several of the less well-characterized flaviviruses have been detected in humans, animals, and mosquitoes across the globe (*2, 3*). Therefore, characterizing these lesser-known viruses is critical to determine whether they have features that portend medically significant future outbreaks.

One such virus is Spondweni virus (SPONV), which is the flavivirus most closely related to ZIKV. SPONV was thought to have been first isolated from a pool of mosquitoes in South Africa in 1955; however, it was later recognized that SPONV was isolated three years earlier from a febrile patient in Nigeria, but because of serological cross-reactivity it was originally thought to be ZIKV (*4–8*). The limited, well-documented human cases describe a clinical presentation similar to ZIKV—most cases result in mild febrile illness, although a subset of these cases document more severe illness including neurological involvement (*5, 8–10*). SPONV is thought to be geographically restricted to Africa. In the era shortly following SPONV’s initial identification, mosquito surveillance, as well as human and animal serosurveys, found evidence of SPONV circulation in 10 sub-Saharan African countries (*5, 11–14*), though serological cross-reactivity with ZIKV and other flaviviruses likely still confounds accurate diagnostics today.

However, in 2016, SPONV RNA was identified in a pool of *Culex quinquefasciatus* mosquitoes in Haiti during routine mosquito surveillance activities (*15*), raising concerns that SPONV was present in the Western Hemisphere and therefore a neglected public health concern. Because human infections with SPONV have historically been sporadic and there have been no known epidemics, neither the disease caused by SPONV nor the mosquito vectors that transmit SPONV have been well-characterized. We recently demonstrated that SPONV can cause significant fetal harm, including demise, comparable to ZIKV in pregnant *Ifnar1*^-/-^ mice. In addition, in pregnant mice treated with an anti-Ifnar1 mAb to transiently abrogate type I interferon signaling prior to SPONV inoculation, we observed infection of the placenta and fetus (*16*), confirming results reported previously (*17*). We also demonstrated that *Aedes aegypti* could efficiently transmit SPONV, whereas *Culex quinquefasciatus* could not (*16*). While these experiments suggested that SPONV may possess features that make it a public health risk, they were performed in immune-compromised mice and therefore may not fully mimic key attributes of human infection, particularly during pregnancy (*18*). A study in the 1950s unwittingly established that rhesus macaques support replication of SPONV (*19*). The inoculum used in those studies was initially thought to be ZIKV but was subsequently shown to be SPONV (*6–8*). The animals apparently developed neutralizing antibodies but no data are provided that describe the virological parameters of the infection.

To assess differences in SPONV replication between macaque species, we infected rhesus (n=4) or cynomolgus (n=5) macaques by subcutaneous inoculation with the South African SPONV isolate SA Ar94. All 4 rhesus macaques were productively infected, with viral load dynamics similar to ZIKV-inoculated controls (n=3). In contrast, all 5 cynomolgus macaques were resistant to SPONV infection. To investigate the breadth of protective immunity induced by a SPONV or ZIKV infection, we also performed a crossover serial challenge experiment in which SPONV-immune animals were rechallenged with the African-lineage ZIKV strain DAK AR 41524 and ZIKV-immune animals were rechallenged with SPONV. Immune responses to SPONV did not provide protection against ZIKV infection. In contrast, immune responses to ZIKV provided protection against SPONV in all animals when rechallenged with a dose of SPONV that productively infected 4/4 naive animals.

## Results

### Cynomolgus macaques are resistant to SPONV infection

Because SPONV is an understudied flavivirus and numerous studies have shown that cynomolgus, rhesus, and pigtail macaques (*Macaca mulatta, M. fascicularis*, and *M. nemestrina*, respectively) are useful platforms to study flavivirus pathogenesis, candidate therapies, and vaccines (reviewed in (*20*)), we sought to characterize SPONV replication dynamics and assess antigenic interactions between SPONV and ZIKV in macaque monkeys. First, n=5 cynomolgus macaques were subcutaneously inoculated with 10^4^ PFU of SPONV strain SA Ar94 (referred to hereafter as SPONV) and n=4 were subcutaneously inoculated with 10^4^ PFU of the African-lineage ZIKV strain DAK AR 41524 (ZIKV-DAK) (Table S1). This dose and route of inoculation was chosen to facilitate comparisons to historical data from ourstudies of ZIKV in macaques (*21–23*). Blood was collected daily for 10 days post inoculation (dpi).

Plasma viral loads were measured by ZIKV- and SPONV-specific quantitative reverse transcription-PCR (RT-qPCR). All four ZIKV-inoculated animals were productively infected with ZIKV, with viral RNA detectable in the plasma by 2 to 4 dpi, viral loads peaking at 10^5^ to 10^6^ viral RNA (vRNA) copies/mL, and duration lasting 4 to 7 days (Fig. 1A). Surprisingly, only 3/5 SPONV-challenged animals had detectable plasma viral loads. In two of these animals, vRNA was detectable in the plasma for 5-6 days with peak viral load only reaching 10^3^ to 10^4^ vRNA copies/mL (Fig. 1A). The third animal had detectable viral loads at only two time points, with a peak vRNA load of 343 copies/mL.

**Figure 1.**
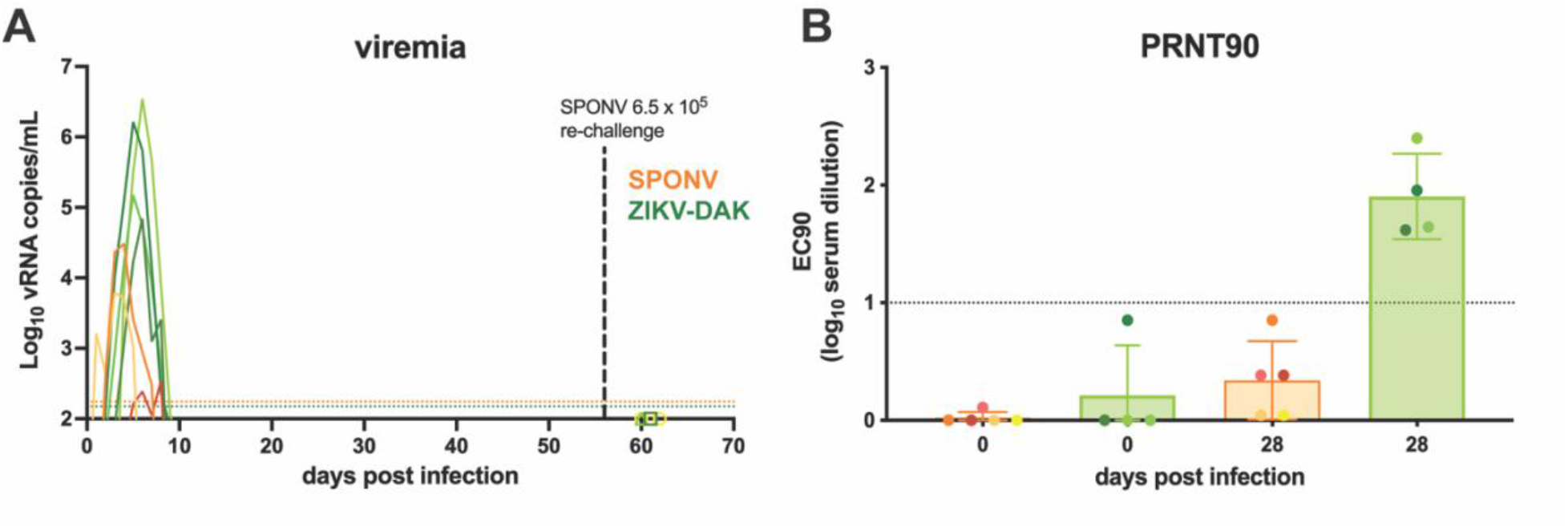
SPONV and ZIKV infection in cynomolgus macaques. (**A)** Plasma viral loads for each of the macaques challenged with 10^4^ PFU of SPONV (orange traces, n = 5) or ZIKV-DAK (green traces, n = 4). All animals were re-challenged with 6.5 X 10^5^ PFU of SPONV 56 days post primary virus challenge. Viral loads were determined using SPONV- and ZIKV-specific QRT-PCR. Only values above the assay’s limit of detection (150 vRNA copies/mL for ZIKV, green dotted line; 175 vRNA copies/mL for SPONV, orange dotted line) are shown. (**B)** PRNT90 titers 0 and 28 days post primary challenge. nAb titers are measured against the same virus stock as used for each animal’s primary challenge (SPONV-challenged sera against SPONV, ZIKV-challenged against ZIKV-DAK). The dotted line represents the PRNT_90_ standard cut-off value of 1:10 dilution determining infection.

Given the limited viral replication in the SPONV-inoculated animals, we next measured serum neutralizing antibody (nAb) responses using plaque reduction neutralization tests (PRNT90). These animals were housed outdoors prior to their arrival at WNPRC, so we cannot define their pathogen exposure history with certainty. However, PRNT90 results confirmed that the SPONV-inoculated animals did not have any pre-existing SPONV antibody response at the time of virus challenge. Similarly, the ZIKV-inoculated animals also were confirmed to be ZIKV naive at the time of challenge (Fig. 1B). We additionally measured nAb titers at 28 dpi, to determine if the SPONV-inoculated animals with detectable viral loads seroconverted. At 28 dpi, all ZIKV-inoculated animals developed robust nAb titers (Fig. 1B), whereas none of the SPONV-challenged animals developed nAb responses to SPONV above the standard 1:20 serum dilution cut-off value that has been traditionally considered diagnostic in the field (*24*)at this time point (Fig. 1B). The SPONV-challenged macaque with the highest viral load and longest duration of detectable viral loads had the highest nAb titer 28 dpi: ∼1:7 (estimated by nonlinear regression). As a result, we cannot robustly conclude that the neutralization response was SPONV-specific and therefore it is unlikely that the transient plasma viral load was the result of productive infection.

While these results suggested varying SPONV susceptibility in cynomolgus macaques, we wanted to exclude the possibility that infection was dose-dependent. Because SPONV-specific nAbs were very low or absent, we s.c. inoculated all nine macaques with 6.5 X 10^5^ PFU SPONV 56 days after the initial virus challenge. This was the highest dose we could administer given the titer of the stock virus. After re-challenge, no animals had detectable SPONV plasma vRNA (Fig. 1A). Although we cannot determine from this experiment whether there was a protective effect from pre-existing immunity in the four animals previously exposed to ZIKV, the consistent results across the dose-range used suggests that cynomolgus macaques are resistant to infection with SPONV.

### SPONV and ZIKV replication in primary cells from cynomolgus and rhesus macaques

We next asked whether primary cells from cynomolgus and rhesus macaques were differentially susceptible to SPONV. Skin fibroblasts have been shown to be permissive to ZIKV infection, and are one of the initial sites of infection for many arboviruses following mosquito-bite inoculation (*25, 26*). We therefore started our characterization of SPONV replication in primary skin fibroblasts derived from adult cynomolgus and rhesus macaques. Fibroblasts were inoculated with an MOI of 0.01 PFU/cell of SPONV or ZIKV-DAK, and infectious virus was quantified via plaque assay from supernatant collected at the time of infection and every 24 hours post-infection (hpi) for the following 5 days (up to 120 hpi). In cynomolgus macaque fibroblasts, the results show a gradual increase in SPONV and ZIKV-DAK titer over time, indicating active replication of both viruses (Fig. 2A). SPONV replication was significantly lower at all timepoints 24-120 hpi compared to ZIKV-DAK in cynomolgus macaque fibroblasts (24-120 hpi: *p* < 0.05, 0 hpi: n.s., unpaired parametric t-test). In rhesus macaque fibroblasts, SPONV and ZIKV-DAK titers also increased over time, indicating that rhesus macaque fibroblasts also support SPONV and ZIKV-DAK replication (Fig. 2B). SPONV replication was also significantly lower than ZIKV-DAK in rhesus macaque fibroblasts at all timepoints 24-120 hpi (24-120 hpi: *p* < 0.01, 0 hpi: n.s., unpaired parametric t-test).

**Figure 2.**
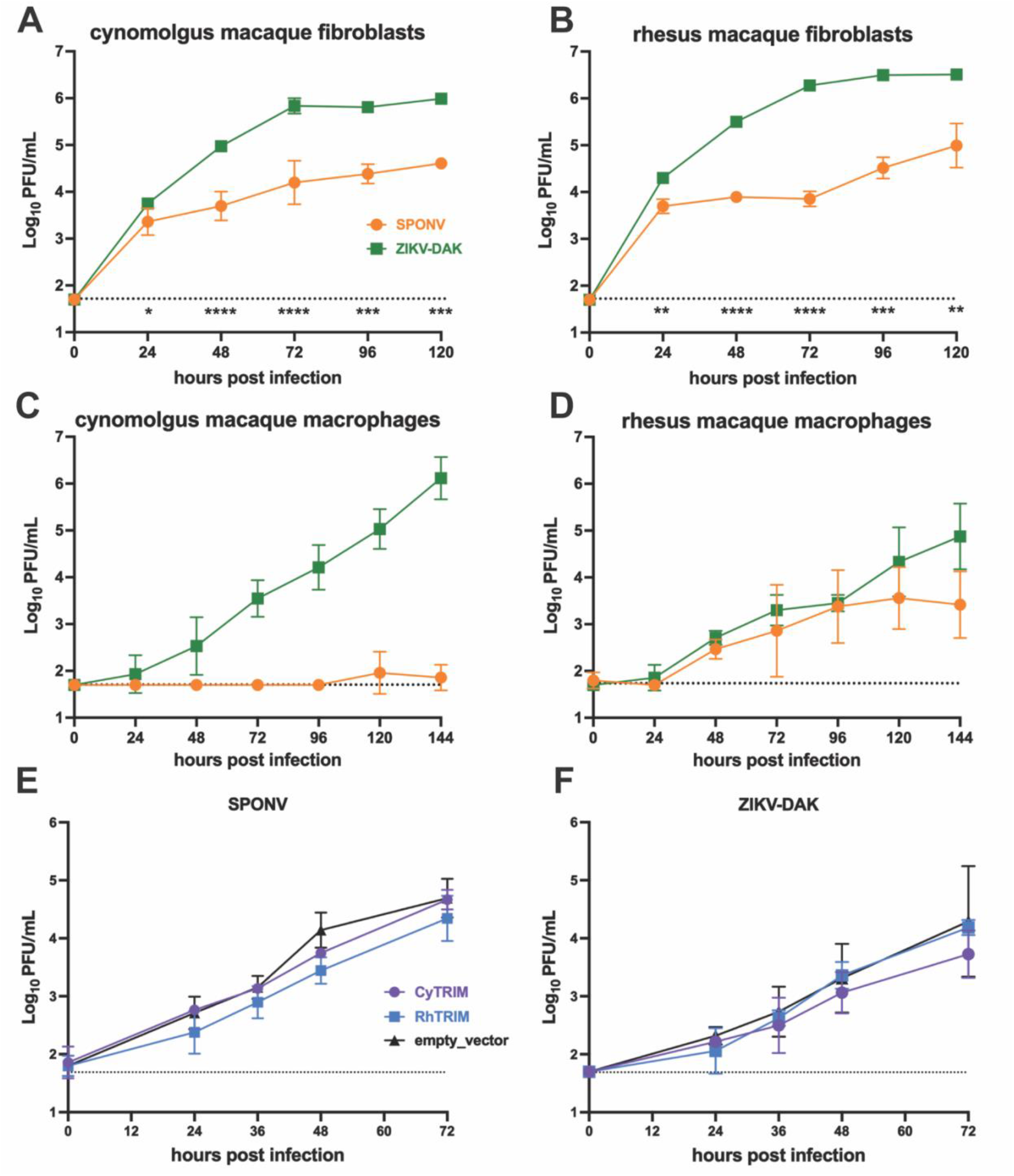
Comparative SPONV and ZIKV replication in vitro. Cynomolgus macaque fibroblasts **(A)**, rhesus macaque fibroblasts **(B)**, cynomolgus macaque macrophages **(C)**, and rhesus macaque macrophages **(D)** were infected with an MOI 0.01 PFU/cell of SPONV (orange) or ZIKV-DAK (green). HEK293 cells expressing cynomolgus (cyTRIM, purple) or rhesus (rhTRIM) TRIM5ɑ, or an empty vector control were infected with an MOI of 0.01 PFU/cell of SPONV **(E)** or ZIKV-DAK **(F)**. Supernatant was collected daily and growth kinetics were assessed by plaque assay. Data presented are from three replicates from one to two independent experiments. Error bars represent standard deviation from the mean. The dotted line indicates the assay limit of detection. Unpaired parametric t-tests with Holm’s-Sidak correction for multiple comparisons were used to test for significance between SPONV and ZIKV-DAK growth kinetics at each timepoint **(A-D)**. *, p < 0.05; **, p < 0.01; ***, p < 0.001; ****, p < 0.0001.

Since both rhesus and cynomolgus macaque fibroblasts supported replication of SPONV and ZIKV, we hypothesized that an innate immune cell could limit SPONV infection in cynomolgus macaques. Macrophages are a key innate immune cell recruited early in response to infection in the skin, are important for ZIKV replication in the skin and blood, and are known to be important for infection of other tissue compartments including the placenta and testes (*27–29*). To test whether cynomolgus macaque macrophages were resistant to SPONV infection, we differentiated macrophages from peripheral blood mononuclear cells (PBMCs) from adult flavivirus-naive cynomolgus and rhesus macaques and measured SPONV and ZIKV replication. We inoculated macrophages from each species at an MOI of 0.01 PFU/cell of SPONV and ZIKV-DAK. Infectious virus was quantified via plaque assay from supernatant collected daily for 6 days. In cynomolgus macaque macrophages, ZIKV-DAK titers increased consistently over time, indicating robust viral replication (Fig. 2C). In contrast, there was no detectable SPONV replication in cynomolgus macaque macrophages at any time point in any of the three replicates, with the exception of 300 PFU/ml in a single replicate at 120 hpi and 150 PFU/ml in a separate replicate at 144 hpi (Fig. 2C). In rhesus macaque macrophages, SPONV and ZIKV-DAK produced similar growth curves that did not significantly differ at any time point (0-144 hpi: *p* > 0.05, multiple unpaired t-tests) (Fig. 2D). Together these data indicate that cynomolgus macaques, but not rhesus macaques, display a resistance mechanism that negatively impacts the infectivity and replicative capacity of SPONV in vitro and in vivo.

### TRIM5ɑ is not the host restriction factor responsible for resistance to SPONV infection in cynomolgus macaques

To begin to understand potential host restriction factors that could be responsible for the replicative barrier for SPONV in cynomolgus macaques, we assessed viral infectivity and replication of ZIKV-DAK and SPONV in vitro using HEK293 cells engineered to stably express cynomolgus macaque (cy) tripartite motif protein 5 (TRIM5ɑ), rhesus macaque (rh) TRIM5ɑ, or an empty vector control.TRIM5ɑ is a well-known HIV host restriction factor that functions in a species-specific manner because of the co-evolution of primates and their ancient lentiviruses (*30–32*). However, recent work has shown that both human and rhesus macaque TRIM5ɑ restrict tick-borne flavivirus replication—with the exception of Powassan virus (POWV)—via proteasomal degradation of the flavivirus protease, NS2B/3 (*33, 34*). A previous study found that a panel of mosquito-borne flaviviruses were not restricted by rhesus or human TRIM5ɑ, but did not investigate the combination of SPONV and cynomolgus macaque TRIM5α (*33*). In our experiments, cyTRIM5ɑ, rhTRIM5ɑ, and cells with an empty vector control supported similar growth for both SPONV and ZIKV-DAK (Fig. 2E and F). These results suggest that TRIM5ɑ is not contributing to the cynomolgus macaque-specific resistance to SPONV.

### Rhesus macaques are susceptible to SPONV infection

The previous experiments establish the ability of SPONV to replicate in multiple cell types isolated from adult rhesus macaques, but replication kinetics in cultured cells cannot capture the complexities of host-pathogen interactions and the generation, distribution, and functional kinetics of innate immune responses to infection within complex tissue environments. To determine whether SPONV infects rhesus macaques, we s.c. inoculated four Indian-origin rhesus macaques (n=2 female, n=2 male) with 10^4^ PFU SPONV and three Indian-origin rhesus macaques (n=1 female, n=2 male) with 10^4^ PFU ZIKV-DAK. This is the same dose and inoculation route used in the cynomolgus macaque experiment described above, as well as in prior ZIKV studies in rhesus macaques conducted by our group (*21, 22, 35, 36*). Following inoculation, all four SPONV-inoculated animals became productively infected, with detectable plasma viral loads starting between 1 and 4 dpi (Fig 3A). SPONV was detectable in plasma for 3 to 6 days, peaking between 2 and 6 dpi at viral loads ranging from 10^4^ to 10^5^ vRNA copies/mL. All ZIKV-inoculated animals were productively infected with ZIKV-DAK (Fig 3A). Peak viral loads in the ZIKV-DAK-challenged cohort ranged from 10^5^ to 10^6^ vRNA copies/mL, which was significantly higher than SPONV (*p* = 0.007, one-way ANOVA with Tukey’s multiple comparisons) (Fig 3B). However, there were no statistically significant differences in area under the curve, duration of viremia, or time to peak viremia between SPONV and ZIKV-DAK (Fig 3B). Additionally, when comparing SPONV replication dynamics to non-pregnant contemporary controls infected with additional ZIKV strains using the same route and dose from (*35*) and (*37*), SPONV replication kinetics did not differ significantly in any parameter tested compared to ZIKV strain PRVABC59, but had significantly lower area under the curve and peak viremia compared to ZIKV strain H/PF/2013 (*35, 37*) (Fig. 3B). Serum neutralizing antibody responses were measured by PRNT90 at 0 and 28 dpi (Fig. 3C), and all animals exhibited robust homotypic nAb responses against the virus used to inoculate each animal. Neutralizing antibody titers generated by the SPONV-inoculated animals against SPONV were not significantly different from those generated by the ZIKV-inoculated animals against ZIKV (SPONV 28 dpi: 2.043 log10; ZIKV 28 dpi: 2.491 log10; *p* = 0.148, unpaired t-test).

**FIG 3:**
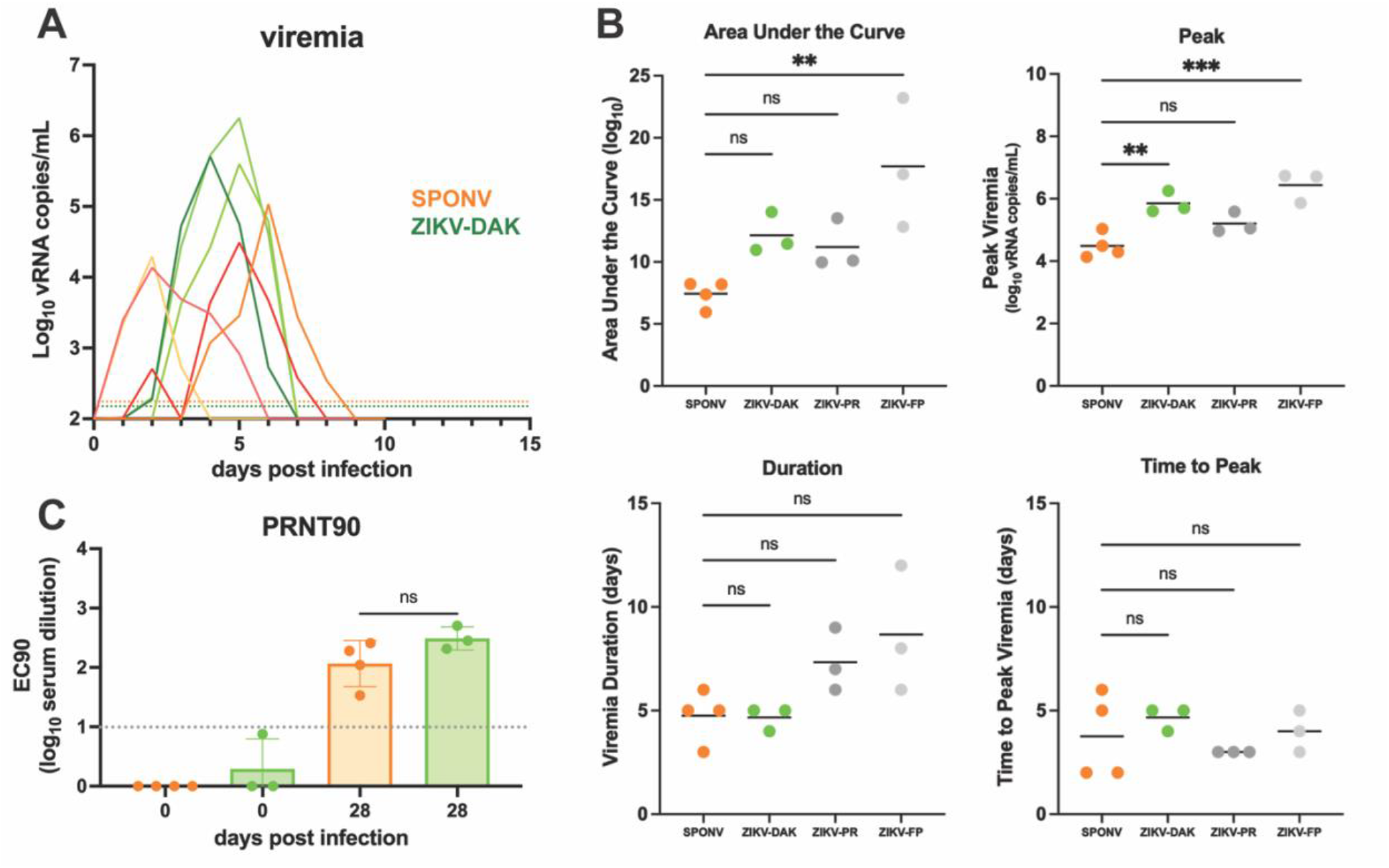
SPONV and ZIKV replication kinetics in rhesus macaques. (**A**) Viral loads were measured from plasma samples from rhesus macaques challenged with 10^4^ PFU of SPONV (n = 4, orange traces) or ZIKV-DAK (n = 3, green traces) using SPONV- or ZIKV-specific QRT-PCR. Only values above the assay’s limit of detection (150 vRNA copies/mL ZIKV, green dotted line; 175 vRNA copies/mL SPONV, orange dotted line) are shown. (**B**) Graphs of the values for the peak, area under the curve, duration, and time to peak viremia. A one-way ANOVA with Tukey’s multiple comparisons test was used for statistical comparison between SPONV and ZIKV-DAK challenged animals, as well as historical data (gray traces) from ZIKV strain PRVABC59 (ZIKV-PR, n = 3) and a French Polynesian strain (ZIKV-FP, n = 3) (\*\*\**p* < 0.0005; \*\**p* < 0.005; \**p* < 0.05; ns, not significant). **(C)** PRNT90 titers from serum collected 0 and 28 dpi. nAb titers are measured against the same virus stock as used for each animal’s primary challenge (SPONV-challenged sera against SPONV, ZIKV-challenged against ZIKV-DAK). An unpaired t-test was used for statistical comparison between SPONV and ZIKV-DAK 28 dpi nAb titers. The dotted line represents the PRNT90 standard cut-off value of 1:10 dilution determining infection.

### Heterologous re-challenge of rhesus macaques results in one-way cross protection between ZIKV and SPONV

Flaviviruses have complex antigenic relationships, in which pre-existing immunity can enhance, attenuate, or have no effect on subsequent infections (*38*). ZIKV and SPONV form a serocomplex and share ∼69% nucleotide identity and ∼75% amino acid identity, so it is conceivable that they may interact antigenically. For reference, the four DENV serotypes— for which it is well-established that pre-existing immunity to one serotype can lead to antibody-dependent enhancement of a secondary infection by a heterologous serotype (*39, 40*)— share 65-70% amino acid identity. It is unknown whether primary infection with SPONV or ZIKV can affect the outcome of subsequent exposure to the heterologous virus. We therefore re-challenged SPONV-immune animals with 1 × 10^4^ PFU of ZIKV-DAK 13 weeks after primary SPONV infection. ZIKV-immune animals were re-challenged with 1 × 10^4^ PFU of SPONV 12 weeks after primary ZIKV-DAK infection.

Upon heterologous re-challenge with ZIKV-DAK, 4/4 SPONV-immune animals became productively infected with ZIKV-DAK (Fig. 4A), but ZIKV-DAK replication dynamics were altered in SPONV-immune animals as compared to in flavivirus-naive animals. When compared to primary infection parameters ZIKV replicated to significantly lower peak plasma viral loads in SPONV-immune animals (*p* = 0.0039, unpaired t-test). ZIKV-DAK area under the curve was also significantly lower in SPONV-immune animals compared to flavivirus-naive animals (*p* = 0.0136, unpaired t-test), but ZIKV-DAK time to peak viral load and viral load duration were not significantly different between SPONV-immune and flavivirus-naive animals (Fig. 4B). Serum neutralizing antibody responses were measured by PRNT50 against SPONV and ZIKV at 0 and 28 days post primary challenge and 0 and 28 days post heterologous rechallenge (91 and 112 days post primary SPONV challenge). For these analyses, PRNT50 titers were more appropriate to compare fine-scale differences in nAb responses in immune animals, due to the higher accuracy of this value within the linear portion of the neutralization curve as compared to PRNT90 values which are preferred for diagnostic identification of flavivirus exposures (*41*). At the time of re-challenge, SPONV-immune animals still had robust neutralizing antibody responses to SPONV as measured by PRNT50 that were not significantly lower than those detected 28 days post primary SPONV infection (2.718 log10 serum dilution vs. 2.178 log10 serum dilution, *p* = 0.312, two-way ANOVA with Tukey’s multiple comparison test). However, these sera did not cross-neutralize ZIKV-DAK (Fig. 4C). At 28 days post secondary ZIKV-challenge, SPONV neutralizing antibody titers were boosted to a significantly higher titer than those detected at 28 days after primary SPONV-challenge (28dp-SPONV: 2.718 log10 serum dilution vs. 28dp-ZIKV: 3.825 log10 serum dilution, *p* = 0.0009, two-way ANOVA with Tukey’s multiple comparisons test). 4/4 animals developed robust ZIKV-specific nAb responses 28 days post secondary ZIKV-challenge (Fig. 4C).

**FIG 4:**
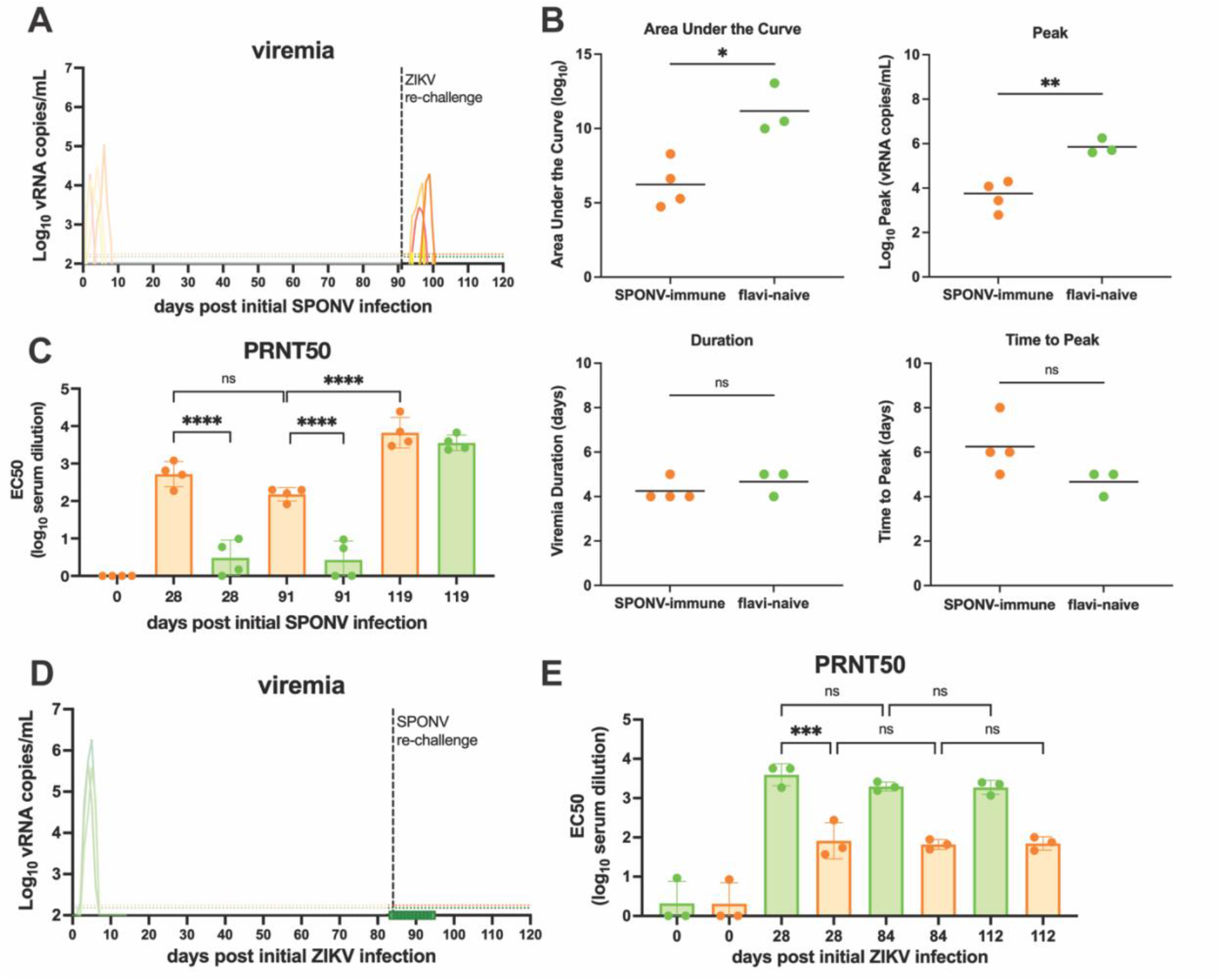
Heterologous re-challenge of SPONV- and ZIKV-immune rhesus macaques. **(A)** Viral loads were measured from plasma samples from rhesus macaques challenged with 10^4^ PFU of ZIKV 91 days post primary SPONV infection (n = 4) using ZIKV-specific QRT-PCR. Only values above the assay’s limit of detection (150 vRNA copies/mL ZIKV green dotted line; 175 vRNA copies/mL SPONV, orange dotted line) are shown. **(B)** Graphs of the values for the area under the curve, peak viremia, viremia duration, and time to peak viremia for ZIKV viremia in SPONV-immune animals (orange) and flavivirus-naive animals (green). An unpaired t-test was used for statistical comparison between groups (**p < 0.005; *p < 0.05; ns, not significant). **(C)** PRNT50 titers from serum collected 0, 28, 91, and 119 days post primary SPONV infection. nAb titers were measured against both SPONV (orange) and ZIKV-DAK (green) at all timepoints. A 2-way ANOVA with multiple comparisons was used for statistical comparison between nAb titers (****p < 0.0001; ns, not significant). **(D)** Viral loads were measured from plasma samples from rhesus macaques challenged with 10^4^ PFU of SPONV 84 days post primary ZIKV infection (n = 3) using SPONV-specific RT-qPCR. **(E)** PRNT50 titers from serum collected 0, 28, 84, and 112 days post primary ZIKV infection. nAb titers were measured against both ZIKV-DAK (green) and SPONV (orange) at all timepoints. A 2-way ANOVA with multiple comparisons was used for statistical comparison between nAb titers (***p < 0.0005; ns, not significant).

Upon heterologous re-challenge with SPONV in ZIKV-immune animals, vRNA was undetectable in plasma at all timepoints through 10 days post re-challenge (Fig. 4D). At the time of secondary SPONV re-challenge, serum nAb titers remained elevated against both SPONV and ZIKV (Fig. 4E). We did not observe an increase in SPONV or ZIKV nAb titers after re-challenge, suggesting that pre-existing ZIKV immunity confers robust protection against SPONV infection (Fig. 4E).

## Discussion

Here we demonstrate that rhesus macaques are susceptible to SPONV infection whereas cynomolgus macaques are resistant. This work thus establishes a nonhuman primate model for SPONV infection. Using this model, we observed one-way cross protection against SPONV in ZIKV-immune animals. This finding is consistent with observations from another study that identified several human cross-reactive mAbs derived from ZIKV- and DENV-infected patients that potently neutralized SPONV in vitro. Passive transfer of some of these mAbs protected mice from lethal SPONV challenge (*17*).

SPONV’s ability to spread and broadly infect new human populations depends in part on susceptible hosts. In ZIKV- and SPONV-endemic regions, people may be infected early in life, developing immunity that protects against subsequent reinfection with the same virus, or limits the pathogenicity of later infection with the heterologous virus. Humans in the Americas had no such protective immunity when ZIKV was introduced, and this may largely explain the scale and scope of the American outbreak. However, if ZIKV immunity provides similarly robust protection against SPONV in humans as we observed in macaques, we speculate that high ZIKV seroprevalence in the Americas [2,8] at the time of SPONV introduction in Haiti in 2016 contributed to limiting SPONV establishment and spread. Importantly, we only assessed cross-protection at a single time point, 12-13 weeks after primary infection, therefore the durability of cross-reactive immunity to SPONV remains uncertain. It is possible that waning of cross-reactive nAb responses occurs more rapidly than homotypic ZIKV immunity so it is unclear how long pre-existing ZIKV immunity will provide robust protection against SPONV (*42, 43*).

Future studies will focus on elucidating the immunological mechanisms that underpin this paradoxical non-reciprocal interaction, because SPONV and ZIKV are not unique in this phenomenon. It is well-established that flaviviruses cross-react. Indeed, cross-reactive antibodies can complicate flavivirus diagnostics, and this feature was initially used to segregate them into distinct serocomplexes (*44, 45*). For example, the sequence of infecting serotypes during serial DENV infection determines whether pre-existing immunity is associated with enhancement or protection (*46, 47*). Likewise, studies of the interaction between ZIKV and DENV suggest that there are asymmetric immune relationships between these viruses as well— DENV infection followed by ZIKV infection has been shown to be cross-protective whereas ZIKV infection followed by DENV-2 infection has been shown to be enhancing in certain scenarios (*48*). Asymmetric immune interactions have also been observed within the tick-borne encephalitis (TBE) serocomplex. Immune sera from tick-borne encephalitis virus (TBEV) vaccinees and sera from infected patients were found to cross-neutralize related viruses within the TBE serocomplex, but did not neutralize POWV, the only North American representative of the TBE serocomplex (*49*). This was posited to be in part due to the lower level of genetic similarity between TBEV and POWV within the envelope (E) glycoprotein EI and EII domains, despite an overall 77% amino acid similarity between TBEV and POWV E protein. For reference, SPONV and ZIKV-DAK share 72% amino acid identity between E proteins with no obvious domain specific differences. A subsequent study testing a POWV mRNA vaccine encoding the prM and E genes found that immune sera from vaccinated mice cross-neutralized a panel of TBE serocomplex viruses—including TBEV—and even protected mice in vivo against the more distantly related Langat virus (*50*). These studies therefore suggest one-way cross-protection between POWV and related TBE serocomplex viruses, however, they do not directly compare cross-protection between these viruses in vivo. Further, it is unclear whether infection-induced versus vaccine-induced immunity generates equivalent amounts of type-specific and cross-reactive antibodies. Many other examples exist of cross-protective immune responses amongst the flaviviruses (*51–53*), however, it is not possible to determine if these responses are asymmetric because the reciprocal sequence of challenges was not performed. Asymmetric immune interactions have also been observed between closely related alphaviruses (*54–56*), and this has been used to formulate hypotheses regarding the lack of alphavirus emergence events, similar to what we postulate may have occurred with SPONV in Haiti.

Although rhesus, cynomolgus, and pigtail macaques are all members of the genus *Macaca*, they have important genotypic and phenotypic differences that can impact the development of animal models (*57, 58*). Because multiple reports (including our own work) previously demonstrated that rhesus, cynomolgus, and pigtail macaques are all susceptible to ZIKV and other flavivirus infections (*20, 59*), we expected that both rhesus and cynomolgus macaques would be susceptible to SPONV infection. However, we observed complete resistance to SPONV infection in cynomolgus macaques. This is similar to what has been described recently for Kyasanur Forest disease virus (KFDV), a tick-borne flavivirus, in rhesus versus pigtail macaques—KFDV is restricted in rhesus macaques but causes moderate to severe disease that recapitulates multiple features of human disease, including hemorrhage in pigtail macaques (*58*). The mechanism(s) underlying resistance to SPONV in cynomolgus macaques is likely multifaceted. However, it was recently shown that the restriction factor TRIM5ɑ robustly inhibited tick-borne flaviviruses but not mosquito-borne flaviviruses (*33*). We examined the ability of cyTRIM5ɑ to restrict SPONV infection because TRIM5ɑ restriction was not universal for the tick-borne flaviviruses (POWV was not restricted by TRIM5ɑ), and restriction for KFDV was primate species-dependent (*58*). Our data suggest that both cyTRIM5ɑ and rhTRIM5ɑ are nonrestrictive for SPONV. Future studies will be needed to elucidate the restriction mechanism(s) controlling this phenotype. However, macaque genetic diversity could confound such studies (*60, 61*). Importantly, our cohort of animals included cynomolgus macaques of both Southeast Asian and Mauritian origin and monkeys from both genetic backgrounds were resistant to SPONV infection. Mauritian-origin cynomolgus macaques have extremely low MHC diversity between animals compared to captive-bred Indian-origin rhesus macaques and cynomolgus macaques from mainland Southeast Asia (*62*). The relatively simple immunogenetics of these animals could be harnessed to identify genes involved in SPONV resistance versus susceptibility. Identifying these factors could provide insight into the evolutionary histories of SPONV and ZIKV and could be vital for understanding the sylvatic reservoirs for SPONV. The natural maintenance cycle of SPONV remains unclear (*6, 10*), but it likely circulates enzootically among unknown vertebrate hosts (presumably nonhuman primates) and is transmitted by arboreal *Aedes* mosquitoes in Africa (*63*).

Although we cannot predict the next major viral epidemic, there is a critical need to improve understanding of understudied viruses, like SPONV, which may also pose a threat. Our study establishes immunocompetent rhesus macaques as a relevant translational model for infection with SPONV. This will enable investigations of immunity, pathogenesis, and medical countermeasures. Critically, it will also enable investigations to define the pathophysiology of SPONV in pregnancy in a model that provides a closer representation of the morphological, developmental, and immune environment at the maternal-fetal interface. The nonreciprocal cross-protection from detectable SPONV infection in ZIKV-immune animals also highlights the increasingly complex heterogeneous immune landscapes that exist in individuals with multiple flavivirus exposures. This has major implications for the flavivirus vaccines that are licensed and commercially available or moving through the clinical pipeline, because most individuals have had multiple exposures to many flaviviruses during their lifetimes. Future studies aimed at characterizing antibody repertories in this system will be valuable to identify the correlates of nonreciprocity between closely-related flaviviruses.

## Materials and Methods

### Ethics statement

This study was approved by the University of Wisconsin-Madison Institutional Animal Care and Use Committee (Animal Care and Use Protocol Number G006256).

### Experimental design

This study was designed to establish the infectivity and replication dynamics of SPONV in a macaque model. A secondary objective was to perform a crossover serial challenge study to better understand the potential for cross-protective immunity between SPONV and ZIKV. Nine cynomolgus macaques (*Macaca fascicularis*) were subcutaneously inoculated with 1 × 10^4^ PFU of SPONV (n = 5) or ZIKV-DAK (n = 4). Cynomolgus macaques (n = 9) were re-challenged with 6.5 X 10^5^ PFU of SPONV 56 days post initial infection. Seven rhesus macaques (*Macaca mulatta*) were subcutaneously inoculated with 1 × 10^4^ PFU of SPONV (n = 4) or ZIKV-DAK (n = 3). 12-13 weeks post-initial infection, rhesus macaques were re-challenged with 1 X 10^4^ PFU of the heterologous virus. Demographic data from the animals from each cohort are provided in (Table S1).

### Care and use of macaques

All macaque monkeys used in this study were cared for by the staff at the Wisconsin National Primate Research Center (WNPRC) in accordance with the regulations and guidelines outlined in the Animal Welfare Act, the Guide for the Care and Use of Laboratory Animals (National Research Council. 2011.), and the recommendations of the Weatherall report (https://royalsociety.org/topics-policy/publications/2006/weatherall-report/). All macaques used in the study were free of Macacine herpesvirus 1, simian retrovirus type D (SRV), simian T-lymphotropic virus type 1 (STLV), and simian immunodeficiency virus. For all procedures (including physical examinations, virus inoculations, and blood collection), animals were anesthetized with an intramuscular dose of ketamine (10 mg/kg). Blood samples were obtained using a Vacutainer system or needle and syringe from the femoral or saphenous vein. Demographic data for animals in each cohort are provided in the table (Table 1) below.

### Cells and viruses

African Green Monkey kidney cells (Vero; ATCC #CCL-81) were maintained in Dulbecco’s modified Eagle medium (DMEM) supplemented with 10% fetal bovine serum (FBS; Hyclone, Logan, UT), 2 mM L-glutamine, 1.5 g/L sodium bicarbonate, 100 U/ml penicillin, 100 μg/ml of streptomycin, and incubated at 37 °C in 5% CO2. *Aedes albopictus* mosquito cells (C6/36; ATCC #CRL-1660) were maintained in DMEM supplemented with 10% fetal bovine serum (FBS; Hyclone, Logan, UT), 2 mM L-glutamine, 1.5 g/L sodium bicarbonate, 100 U/ml penicillin, 100 μg/ml of streptomycin, and incubated at 28 °C in 5% CO2. Human embryonic kidney cells (HEK-293; ATCC #CRL-1573) were maintained in DMEM supplemented with DMEM supplemented with 10% FBS, 2mM L-glutamine, 1.5% g/L sodium bicarbonate, 100U/ml penicillin, 100 μg/ml of streptomycin, and incubated at 37 °C in 5% CO2. The cell lines were obtained from the American Type Culture Collection, were not further authenticated, and were tested and confirmed negative for mycoplasma.

#### Primary cell lines

Fibroblasts were differentiated from skin punch biopsies from adult rhesus and cynomolgus macaques. Fibroblasts were confirmed Herpes B and mycoplasma negative. Fibroblasts were maintained in in Dulbecco’s modified Eagle medium (DMEM) supplemented with 20% fetal bovine serum (FBS; Hyclone, Logan, UT), 2 mM L-glutamine, 1.5 g/L sodium bicarbonate, 100 U/ml penicillin, 100 μg/ml of streptomycin, 1% MEM 100X non-essential amino acids, and incubated at 37 °C in 5% CO2.

Macrophages were derived from peripheral blood mononuclear cells (PBMCs) from flavivirus-naive adult rhesus and cynomolgus macaques. Macrophages were differentiated as previously described (*64*). At 4-5 days post treatment of adherent cells with supplemented media containing M-CSF (Peprotech) and IL-1β (Peprotech), cells were detached with a cell scraper and replated in twelve-well plates to conduct virus growth curves. A subset of cynomolgus macaque cells were processed for flow cytometry analysis to confirm macrophage differentiation (fig. S1).

ZIKV strain DAK AR 41524 (ZIKV-DAK; GenBank:<u>KY348860</u>) was originally isolated from *Aedes africanus* mosquitoes in Senegal in 1984, with a round of amplification on *Aedes pseudocutellaris* cells, followed by amplification on C6/36 cells, followed by two rounds of amplification on Vero cells, was obtained from BEI Resources (Manassas, VA). SPONV strain SA Ar94 (GenBank:<u>KX227370</u>) was originally isolated from a *Mansonia uniformis* mosquito in Lake Simbu, Natal, South Africa in 1955, with five rounds of amplification with unknown culture conditions followed by a single round of amplification on Vero cells. Virus stocks were prepared by inoculation onto a confluent monolayer of C6/36 mosquito cells. We deep sequenced our virus stocks to verify the expected origin. The SPONV and ZIKV-DAK stocks matched the GenBank sequences (<u>KY348860</u>, <u>KX227370</u>, respectively) of the parental viruses; but a variant at site 3710 in the ZIKV-DAK stock encodes a nonsynonymous change (A to V) in NS2A.

### Plaque assay

All ZIKV and SPONV screens from growth curves and titrations for virus quantification from virus stocks were completed by plaque assay on Vero cell cultures. Duplicate wells were infected with 0.1 ml aliquots from serial 10-fold dilutions in growth media and virus was adsorbed for 1 h. Following incubation, the inoculum was removed, and monolayers were overlaid with 3 ml containing a 1:1 mixture of 1.2% oxoid agar and 2X DMEM (Gibco, Carlsbad, CA) with 10% (vol/vol) FBS and 2% (vol/vol) penicillin/streptomycin. Cells were incubated at 37°C in 5% CO2 for four days for plaque development for ZIKV and five days for SPONV. Cell monolayers then were stained with 3 ml of overlay containing a 1:1 mixture of 1.2% oxoid agar and 2X DMEM with 2% (vol/vol) FBS, 2% (vol/vol) penicillin/streptomycin, and 0.33% neutral red (Gibco). Cells were incubated overnight at 37 °C and plaques were counted.

### Inoculations

Inocula were prepared from the viral stocks described above. The stocks were thawed, diluted in PBS to 1 × 10^4^ PFU/ml for all inocula except for the re-challenge of cynomolgus macaques for which stocks were diluted to 6.5 × 10^5^ PFU/ml. Diluted inocula was then loaded into a 3-ml syringe that was kept on ice until challenge. Animals were anesthetized as described above, and 1 ml of the inoculum was delivered subcutaneously over the cranial dorsum. Animals were monitored closely following inoculation for any signs of an adverse reaction.

### Viral RNA isolation

Viral RNA was extracted from plasma using the Viral Total Nucleic Acid Kit (Promega, Madison, WI) on a Maxwell 48 RSC instrument (Promega, Madison, WI). RNA was then quantified using quantitative RT-PCR. Viral load data from plasma are expressed as vRNA copies/mL.

### Quantitative reverse transcription PCR (QRT-PCR)

vRNA isolated from plasma samples was quantified by quantitative reverse transcription-PCR (RT-qPCR) as described previously (*65*). The SPONV, primer and probe sequences are as follows; forward primer: 5’-GGCATACAGGAGCCACATCAAAC-3’, reverse primer: 5’-TGCGTGGGCTTCTCTGAA-3’ and probe; 5’-6-carboxyfluorescein-CATCACTGGAACAAYAAGGAGGCGCTGG-BHQ1-3’. The RT-PCR was performed using the SuperScript III Platinum One-Step Quantitative RT-PCR system (Invitrogen, Carlsbad, CA) or Taqman Fast Virus 1-step master mix (Applied Biosystems, Foster City, CA) on a LightCycler 96 or LightCycler 480 instrument (Roche Diagnostics, Indianapolis, IN). The viral RNA concentration was determined by interpolation onto an internal standard curve composed of seven 10-fold serial dilutions of a synthetic ZIKV or SPONV RNA fragment. The ZIKV RNA fragment is based on a ZIKV strain derived from French Polynesia that shares >99% identity at the nucleotide level with the African lineage strain used in the infections described in this report. The SPONV RNA fragment is based on the same SPONV strain derived from South Africa used in the experiments in this manuscript. Lower limit of detection (LLOD) for the ZIKV RT-qPCR assay is 150 vRNA copies/mL. LLOD for the SPONV RT-qPCR assay is 175 vRNA copies/mL. LLOD of assays is defined as the cut-off for which plasma viral loads are true positive with 95% confidence.

### Plaque reduction neutralization test (PRNT)

Macaque serum was isolated from whole blood on the same day it was collected by using a serum separator tube (SST). The SST was centrifuged for a minimum of 20 min at 1,400 × g, and the serum layer was removed, placed in a 15-ml conical tube, and centrifuged for 8 min at 670 × g to remove any additional cells. Serum was screened for ZIKV and SPONV neutralizing antibodies by plaque reduction neutralization test (PRNT) on Vero cells as described in reference (*66*) against ZIKV and SPONV. The neutralization assay was performed with the same virus stocks that were used for the challenge. Neutralization curves were generated using GraphPad Prism 8 software. The resulting data were analyzed by nonlinear regression to estimate the dilution of serum required to inhibit 90% of 50% of infection.

### In vitro viral replication

Six-well plates containing confluent monolayers of rhesus or cynomolgus macaque fibroblasts, were inoculated with virus (SPONV or ZIKV-DAK), in triplicate at a multiplicity of infection of 0.01 PFU/cell. After one hour of adsorption at 37°C, inoculum was removed and the cultures were washed three times. Fresh media were added and the fibroblast cultures were incubated for 5 days at 37°C with aliquots removed every 24 hours and stored at -80C. Viral titers at each time point were determined by plaque titration on Vero cells. The same methodology and multiplicity of infection was followed for quantifying in vitro viral replication of SPONV and ZIKV-DAK in rhesus and cynomolgus macaque macrophages, and TRIM5ɑ expressing HEK-293 cells. For macrophage growth curves, 12-well plates were used to achieve a confluent monolayer and samples were collected for an additional 2 days. For TRIM5ɑ expressing HEK-293 cells, supernatant was additionally collected 36 HPI.

### Generation of TRIM5ɑ expressing cells

HEK293 cells stably expressing TRIM5ɑ were generated as previously described in (*33*). Plasmid DNA encoding rhesus macaque (GenBank: EF113914.1) and cynomolgus macaque (GenBank: AB210052.1) TRIM5ɑ open reading frames were ordered from Twist Biosciences and subcloned into MIG1R-derived simple retroviral transduction vectors (*67*) encoding a blasticidin resistance gene downstream of an internal ribosome entry site. To generate retrovirus for transducing TRIM5ɑ expressing vectors, pre-adhered HEK293 cells in 6-well plates were transfected with 1μg vector plasmid, 1μg pMD.Gag/GagPol (*68*) plasmid, and 200ng VSV-G (*69*). Media was replaced at 24 hours post-transfection. Virus-containing supernatant was harvested at 48 hours post-transfection, 0.45μm syringe-filtered, and stored at -20 degrees. To generate stable cells, HEK293 cells were seeded into plates and allowed to adhere overnight and transducing viral supernatant with 10μg/mL polybrene was added to each well. Transduced cells were selected at 48 hours post-transduction with 8μg/mL Blasticidin S (GoldBio, #B-800-100), expanded, and maintained in culture in the presence of drug. Rhesus and cynomolgus TRIM5ɑ restriction activity against HIV-1 was confirmed by single cycle infectivity assay (fig. S2). Briefly, equivalent numbers HEK293 cells transduced to express rhesus or cynomolgus TRIM5ɑ (as well as vector transduced cells) were infected with single cycle HIV-1 virus (NL4-3 Env-Vpr-Nef-mCherry=reporter) (*70*) or murine leukemia virus pseudovirus (MLV gag/gagpol virus-like particle packaging an mCherry expressing genomic RNA) (*67*) pseudo-typed with VSV-G at multiple MOIs. After 48 hours, percent of target cells expressing mCherry (successfully infected) was determined by flow cytometry (BD FACSCanto II).

### Statistical analyses

All statistical analyses were performed using GraphPad Prism 9. For statistical analysis of virus growth curves, unpaired nonparametric t-tests with Holm’s-Sidak correction for multiple comparisons were used to compare SPONV and ZIKV titers at each timepoint. Ordinary one-way ANOVA with Tukey’s multiple comparisons were used to statistically compare differences in area under the curve, peak viremia, time to peak viremia, and viremia duration between macaques infected with SPONV and those infected with ZIKV-DAK, as well as historical viremia data of rhesus macaques infected with ZIKV-PR and ZIKV-FP. The LLOD for SPONV (175 vRNA copies/mL) was used as the baseline for AUC comparisons between virus groups. Unpaired nonparametric t-tests were used to compare area under the curve, peak viremia, time to peak viremia, and viremia duration between flavivirus naive macaques infected with ZIKV-DAK and SPONV-immune macaques infected with ZIKV-DAK.

## Acknowledgments

We thank the Veterinary Services, Colony Management, Scientific Protocol Implementation, and the Pathology Services staff at the Wisconsin National Primate Research Center (WNPRC) for their contributions to this study.

## Author contributions

ASJ: Conceptualization, Validation, Formal analysis, Investigation, Writing - original draft, Writing - review & editing, Visualization. CMC: Conceptualization, Investigation, Writing - review & editing. AMW: Investigation. Writing - review & editing. MIB: Investigation. SLR: Investigation, Writing - review & editing. AR: Investigation. ME: Investigation. EP: Investigation. SC: Supervision. AB: Investigation. JTB: Investigation, Writing - review & editing. JTG: Investigation, Writing - review & editing. TSF: Supervision, Writing - review & editing. RAL: Supervision. TCF: Methodology, Resources, Data curation, Writing: review & editing, Supervision, Funding acquisition. MTA: Conceptualization, Methodology, Formal analysis, Investigation, Resources, Writing - original draft, Writing - review & editing, Supervision, Funding acquisition.

## Data and materials availability

All data needed to evaluate the conclusions in the paper are present in the paper and/or the Supplementary Materials.

