## Supplementary material for "Primary infection with Zika virus provides one-way heterologous protection against Spondweni virus infection in rhesus macaques": SPONV_NHP_SuppMat_Jaeger2022.html

Supporting Online Material for


Supplementary Materials
for

**Primary infection with
Zika virus provides one-way heterologous protection against Spondweni
virus infection in rhesus macaques**

Anna Jaeger *et
al.*

\*Corresponding author.


**This PDF file includes:**

Figs.
S1 to S2

Table
S1

  


Fig.
S1. Flow cytometry analysis of PBMC-derived
macrophages. Surface marker expression of PBMC-derived
macrophages from cynomolgus macaques were analyzed by flow cytometry to confirm
successful differentiation. Cells had high surface expression levels of CD14,
CD16, HLA-DR, CD206, and CD86; the dendritic cell marker CD123 was not
expressed. Abbreviations: FSC-H (forward scatter-height), SSC-A (side
scatter-area).

 


**Fig.
S2. Macaque TRIM5a inhibits HIV, but not MLV.** HEK293 cells transduced as indicated (vector,
rhTRIM5a-HA, cyTRIM5a-HA) were infected with single cycle HIV-1 or murine
leukemia virus (MLV) mCherry reporter viruses at
approximate MOI/cell of 1.0 and 0.1. After 48 hours, % of cells that were mCherry positive was determined by flow cytometry. HIV-1
was restricted by both rhesus and cynomolgus TRIM5-alpha whereas MLV was
unaffected.
